# Prophylactic protection against respiratory viruses conferred by a prototype live attenuated influenza virus vaccine

**DOI:** 10.1101/2021.04.28.441797

**Authors:** Raveen Rathnasinghe, Mirella Salvatore, Hongyong Zheng, Sonia Jangra, Thomas Kehrer, Ignacio Mena, Michael Schotsaert, Thomas Muster, Peter Palese, Adolfo García-Sastre

**Affiliations:** Department of Microbiology, Icahn School of Medicine at Mount Sinai, New York, NY 10029, USA; Graduate School of Biomedical Sciences, Icahn School of Medicine at Mount Sinai, New York, NY 10029, USA; Global Health and Emerging Pathogens Institute, Icahn School of Medicine at Mount Sinai, New York, NY 10029, USA; Department of Medicine, Weill Cornell Medical College, New York, NY, USA; Department of Dermatology, University of Vienna Medical School, 1090 Wien, Austria; Department of Medicine, Division of Infectious Diseases, Icahn School of Medicine at Mount Sinai, New York, NY 10029, USA; The Tisch Cancer Institute, Icahn School of Medicine at Mount Sinai, New York, NY 10029, USA

**Keywords:** Type I IFN, Antiviral Therapy, NS1 protein, Influenza A, Interferon Antagonists, SARS-CoV-2

## Abstract

The influenza A non-structural protein 1 (NS1) is known for its ability to hinder the synthesis of type I interferon (IFN) during viral infection. Influenza viruses lacking NS1 (ΔNS1) are under clinical development as live attenuated human influenza virus vaccines and induce potent influenza virus-specific humoral and cellular adaptive immune responses. Attenuation of ΔNS1 influenza viruses is due to their high IFN inducing properties, that limit their replication in vivo. This study demonstrates that pre-treatment with a ΔNS1 virus results in an immediate antiviral state which prevents subsequent replication of homologous and heterologous viruses, preventing disease from virus respiratory pathogens, including SARS-CoV-2. Our studies suggest that ΔNS1 influenza viruses could be used for the prophylaxis of influenza, SARS-CoV-2 and other human respiratory viral infections, and that an influenza virus vaccine based on ΔNS1 live attenuated viruses would confer broad protection against influenza virus infection from the moment of administration, first by non-specific innate immune induction, followed by specific adaptive immunity.

## Introduction

The type I interferon (IFN) response resulting from invading viral pathogens is considered as one of the first lines of antiviral defence mechanisms in higher organisms. The latter process takes place upon the detection of the pathogen associated molecular patterns (PAMPS) by the host pattern recognition receptors (PRRs). Secretion of interferons takes place in both paracrine and autocrine signalling mechanisms, mediated by the canonical JAK/STAT signal transduction pathway along with the transcriptional activation of a particular set of host genes as well as their corresponding promotors defined as IFN-stimulated response elements (ISREs)^1^. Subsequent activation of the downstream interferon stimulated genes (ISGs) lead to the transcriptional induction of a plethora of antiviral proteins, including dsRNA-activated protein kinase (PKR) leading to a halt of protein translation, dsRNA-activated oligoadenylate synthetases (OAS) which facilitate the degradation of RNA by activating RNAse L and Mx proteins which essentially sequester incoming viral components such as nucleocapsids^2, 3^. Many studies have demonstrated that viruses have evolved to encode numerous mechanisms to prevent the host IFN-mediated antiviral response at different stages^4^. Viral non-structural proteins such as those of Toscana virus, dengue and HPV can sequester host factors to inhibit type I IFN response^5,6,7^, while viruses such as vaccinia, adeno and Ebola viruses secrete soluble ligands^7,8^, or encode miRNAs^9, 10^ and other proteins to confer immune-evasion.

The influenza A virus (IAV) non-structural protein 1 (NS1) facilitates several functions ranging from inhibition of host mRNA polyadenylation and subsequent inhibition of their nuclear export as well as inhibition of pre-mRNA splicing^11, 12^. A growing body of evidence to date has indicated that influenza NS1 protein has IFN antagonistic activity. It was initially shown that a recombinant influenza A virus that lacks the NS1 protein(ΔNS1) grew to a titer similar to that of WT virus in IFN deficient systems, albeit being markedly attenuated in IFN competent hosts^13^. This attenuated phenotype can be explained by the inability of the virus to prevent NS1 mediated IFN inhibition. The NS1 protein has been shown to bind to TRIM25 whereby the ubiquitination of the viral RNA sensor RIG-I is inhibited, which eventually results in the inhibition of IFN induction^14,15^. NS1 has also been shown to prevent IFN production by sequestering the cellular cleavage and polyadenylation specificity factor 30 (CPSF30) in order to halt the processing of host pre-mRNAs, resulting in accumulation of pre-mRNAs in the nucleus as well as the halt of cellular mRNA export to the cytoplasm^16^. This subsequently results in the inhibition of host protein production, including IFNs and proteins encoded by IFN inducible genes ^17,18^ NS1 has also been shown to inhibit the antiviral activity of several IFN-stimulated genes, such as the 2’-5’-oligo A synthase (OAS)^19^.

Consistent with its function, deletion of NS1 in recombinant IAV results in a live attenuated and highly immunogenic IAV. As a result, IAV with impaired NS1 function are currently used as vaccines against swine influenza in pigs^20^ and they are under clinical consideration as live attenuated human influenza virus vaccines^21–23^.

Based on the growing body of evidence showing the IFN antagonistic properties of IAV NS1, we investigated the ability of the ΔNS1 viruses to induce an immediate IFN response *in vivo* along with the biological antiviral consequences mediated by the type I IFN induction. Our results demonstrate that the ΔNS1 virus is an efficient inducer of IFN with antiviral properties in both mice and embryonated eggs. Our data indicates the suitability of ΔNS1 virus as a prophylactic agent to induce immediate mucosal antiviral responses with the aim of preventing acute respiratory infections caused by IFN sensitive viruses. ΔNS1 influenza viruses can provide first innate antiviral protection, followed by adaptive specific IAV protection.

## Results

### Recombinant influenza A virus lacking the NS1 gene (ΔNS1) induces higher levels of interferon than wild type viruses in embryonated chicken eggs

Previously, we demonstrated that tissue culture-based infections by ΔNS1 viruses induced the transactivation of an ISRE-containing reporter gene^13^, indicating that infection by ΔNS1 viruses induces higher levels of IFN in comparison to its wild type counterparts. To test whether ΔNS1induces IFN in 10-day old embryonated-chicken eggs, eggs were treated with 10^3^ PFU of ΔNS1 or PR8-WT influenza viruses. Subsequently, the allantoic fluids were harvested 18 hours post treatment to measure the levels of IFN by determining the highest dilution that inhibited the cytopathic effect mediated by vesicular stomatitis virus (VSV) in chicken embryo fibroblast (CEF) cells. As indicated in the Supplementary table 1, four hundred Uml^−1^ of IFN were detected in the allantoic fluid of eggs infected by ΔNS1 virus. However, allantoic fluids derived from WT-PR8 or mock infections indicated undetectable levels of IFN (<16 Uml^−1^).

### Pre-treatment with ΔNS1 influenza virus inhibits wild-type viral replication in embryonated chicken eggs

We speculated that the ability of the ΔNS1 virus on inducing high titers of IFN in eggs facilitates an antiviral state that may prevent the replication of wild-type IAV. To evaluate this, increasing amounts of ΔNS1 virus were inoculated into eggs and eight hours post-treatment, the eggs were challenged with wild-type A/WSN/33 (WSN-WT) virus with a dose of 10^3^ PFU. Two days post incubation extracted allantoic fluids were titrated via plaque assays. WSN viral titers decreased with ΔNS1 in a dose dependent manner. While the untreated allantoic fluids supported the growth of WSN virus to an approximate titer of 10^8^ PFUml^−1^, administration of a dose as little as 2×10^4^ PFUml^−1^ of ΔNS1 prevented the replication of WSN virus (less than 10^2^ PFUml^−1^ of WSN were obtained in eggs). The titer of WSN virus was reduced by one log, by pre-treating allantoic fluids with as little as 2 PFU of ΔNS1 (Figure 1A).

**Figure 1.**
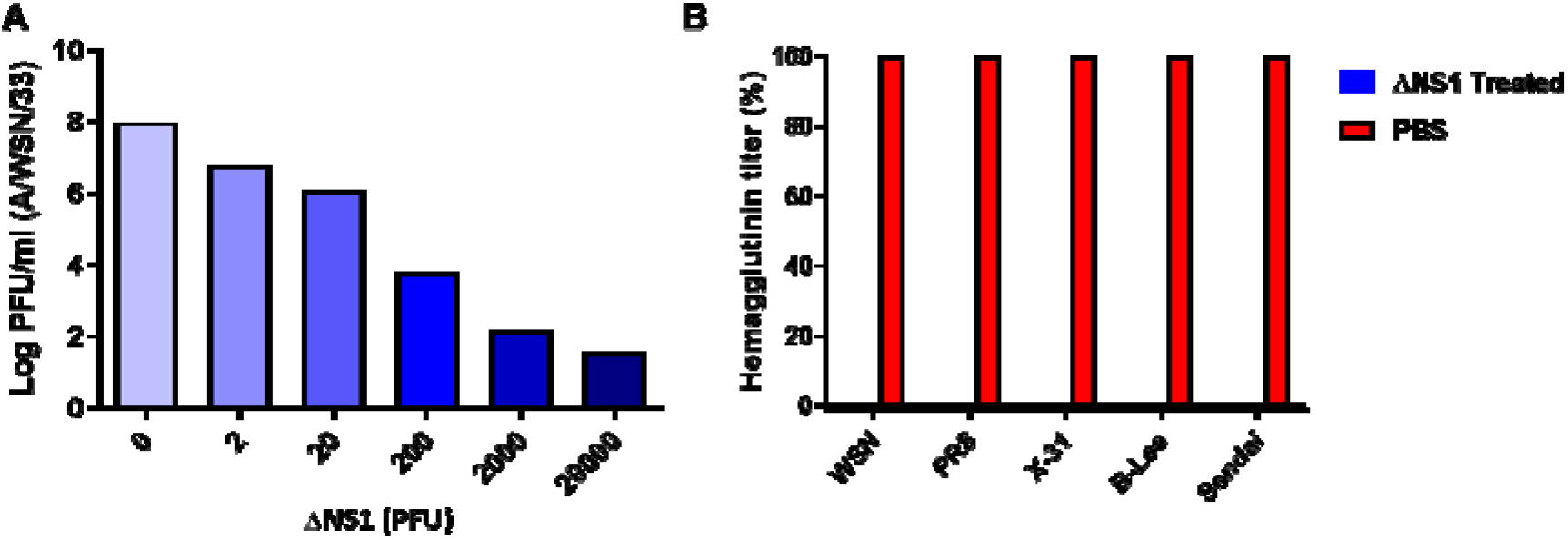
Pre-incubation with ΔNS1 virus inhibits viral replication in embryonated chicken eggs. **(A)** 10-day-old embryonated chicken eggs (n=2 per group) were inoculated with varying amounts of (PFU) of ΔNS1 virus in the allantoic cavity. Eight hours post infection at 37°C, eggs were re-infected with 10^4^ PFU of WT A/WSN/33 influenza virus and incubated at 37°C for 40 hours. Allantoic fluids were then titrated by plaque assay MDBK cells. **(B)** 10-day-old embryonated chicken eggs (n=2 per group) were inoculated with 2×10^4^ PFU of ΔNS1 virus or PBS (Untreated). 8 hours post inoculation at 37°C, the eggs were re-infected with 10^3^ PFU of A/WSN/33 (WSN/H1N1), A/PR/8 (PR8/H1N1), A/X-31 (X-31/H3N2), B/Lee/40 (B-Lee influenza B) or Sendai Virus (Sendai). B-Lee infected eggs were incubated at 35°C for additional 40 h. All other eggs were incubated at 37°C for additional 40 h. Virus present in the allantoic fluid was titrated by hemagglutination assays. Maximum hemagglutination titers (100%) for each individual virus were 2048 (PR8), 1024 (X-31), 256 (B-Lee), 512 (Sendai)

Interestingly, treatment using ΔNS1 virus further inhibited the replication of other viruses, as depicted in figure 1B. Relative HA titers were obtained from eggs treated with 2×10^4^ PFUml^−1^ of ΔNS1 virus followed by subsequent infection with wild-type Influenza A H1N1 strains WSN and PR8, H3N2 strain X-31, influenza B virus or Sendai virus (SeV; a paramyxovirus). In all cases, pre-treatment with ΔNS1 resulted in a two-log reduction of wild-type viral HA titers.

### Severe disease and death caused by infection with the highly virulent PR8 virus (hvPR8) in A2G mice can be alleviated by ΔNS1 pre-treatment

In order to assess whether or not the administration of ΔNS1 virus inhibits replication of influenza viruses in mice, an inbred mouse strain that is homozygous for the gene which codes for the IFN induced full-length *Mx1* protein, defined as C57BL/6-A2G (abbreviated as A2G) mice were used for this part of the study^24, 25^. Previous studies have concluded that IFN administration was ineffective in preventing IAV replication in laboratory mice lacking a functional *Mx1* gene^26^. In contrast, A2G mice which were administered IFN remained alive upon infection with the highly virulent hvPR8 IAV strain^27^. The presence of a functional *Mx1* gene in A2G mice better mirrors the human situation, as *Mx1* gene deficiencies in humans are rare. Here, A2G mice were intranasally infected with a dose of 5×10^5^ PFUml^−1^ of ΔNS1 virus or PBS at −24, −8, +3, +24 and +48 hours. Mice were challenged at time 0 intranasally with 5×10^6^ PFU of hvPR8 virus. Mice treated with ΔNS1 virus were protected from hvPR8 virus as measured by weight loss and death while the PBS treated mice succumbed to death (Figure 2A).

**Figure 2.**
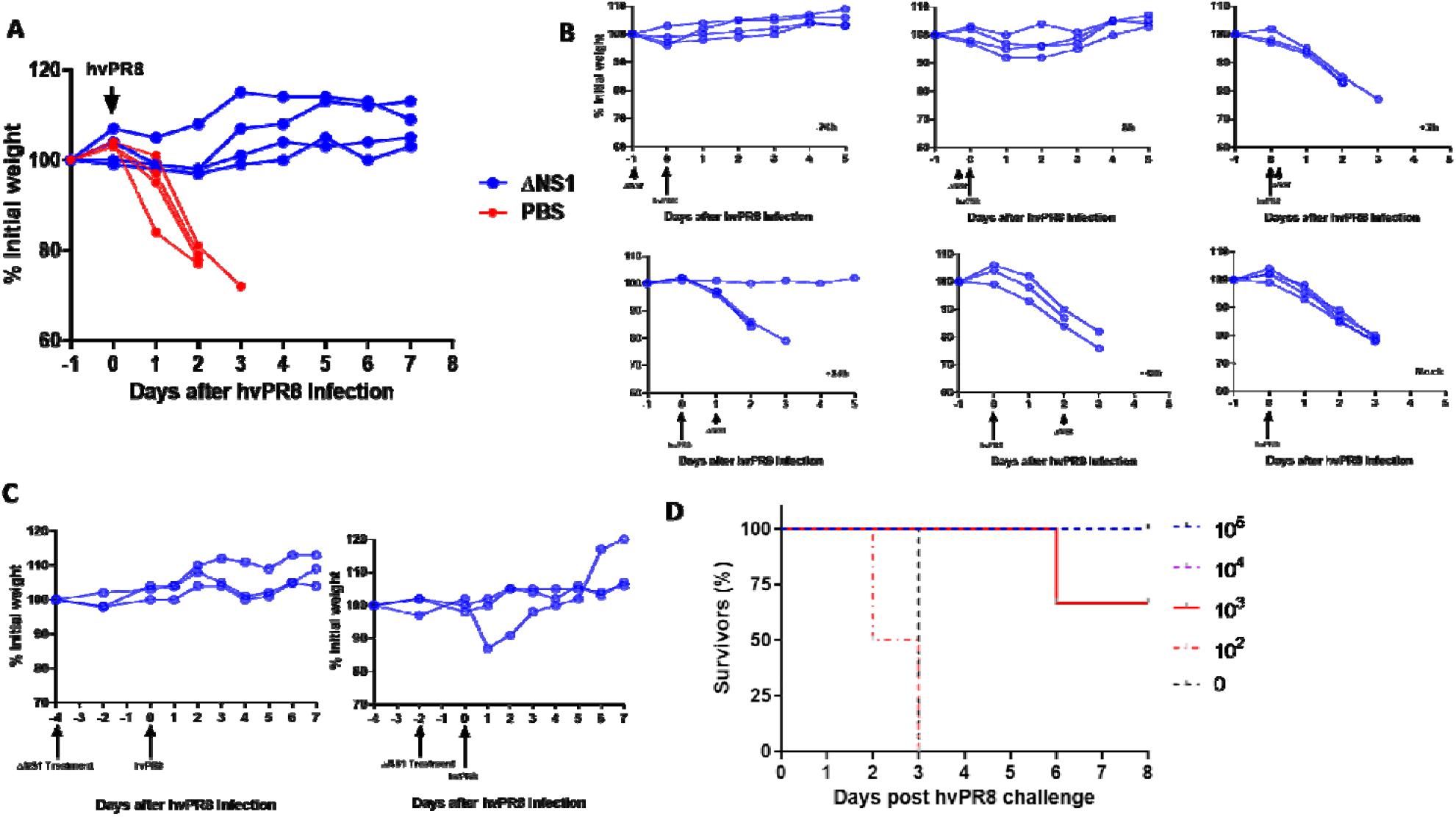
A single dose of ΔNS1 virus protects A2G mice against lethal infection by highly virulent hvPR8 influenza virus when given prior to virus challenge. **(A) Treatment with ΔNS1 virus protects A2G mice against lethal infection by highly virulent hvPR8 influenza virus**. Eight 6-week old A2G mice were intranasally infected with 5×10^6^ PFU of highly virulent A/PR/8/34 (hvPR8) influenza virus. Half of the mice received a total of five intranasal treatments with 5×10^5^ PFU of ΔNS1 virus at the following times with respect to the hvPR8 infection: −24 h, −8 h, +3 h, +24 h ad 48 h. The remaining four mice were treated with PBS and the bodyweight changes and survival was monitored. **(B) A single dose of ΔNS1 virus protects A2G mice against lethal infection by highly virulent hvPR8 influenza virus when given prior to hvPR8 virus challenge**. Groups of three A2G mice each were mock-treated or treated intranasally with 5×10^5^ PFU of ΔNS1 at time points −24 h, −8 h, +3h, +24h, +48h relative to the intranasal infection by 5×10^6^ hvPR8 influenza virus. **(C) A single dose of ΔNS1 virus protects A2G mice against lethal infection by highly virulent hvPR8 influenza virus when given two and four days prior to hvPR8 virus administration** Groups of three A2G mice were intranasally treated with 5×10^5^ PFU of ΔNS1 virus four days or two days before infection by 5×10^6^ hvPR8 influenza virus. Bodyweight changes and survival was monitored. All data points are from individual mice. **(D) Determination of the minimal effective therapeutic dose of ΔNS1 to prevent lethal hvPR8 virus infection in A2G mice**. Groups of three A2G mice were intranasally infected with 10^5^, 10^4^ or 10^3^ PFU ΔNS1 influenza virus. Additionally, groups of two A2G mice were intranasally challenged with 10^2^ of ΔNS1 virus or PBS. 24 hours post inoculation, mice were challenged with by 5×10^6^ hvPR8 influenza virus. The percentage of mice surviving the challenge is represented.

Subsequently, we examined whether all five ΔNS1 treatments were essential for the protective effect against hvPR8 infection in mice. Hence, a single dose of 5×10^6^ PFU of ΔNS1 virus was given at various time points relative to the infection with hvPR8. Data indicated (Figure 2B) that pre-treatment (hours 24 or 8 before hvPR8 challenge) but not post treatment (even 3 hours post hvPR8 challenge) of ΔNS1 resulted in the prevention of weight loss disease and subsequent death. Additionally, ΔNS1 virus administered two or four days prior to hvPR8 challenge completely protected mice from disease (Figure 2C).

Next, to obtain the effective dose 50 (ED_50_) of ΔNS1 virus to mediate protection against disease from hvPR8 infection, 2×10^5^, 2×10^4^, 2×10^3^or 2×10^2^ doses of ΔNS1 virus were intranasally administered to A2G mice 24 hours prior to hvPR8 challenge. As shown in Figure 2D, the ED_50_ of the ΔNS1 virus which conferred protection in A2G mice against hvPR8-induced death was approximately 10^3^ PFU.

### Induction of *Mx1* specific mRNA in mice treated with ΔNS1 virus

To investigate whether ΔNS1 infection in mice resulted in induction of the *Mx1* gene, an RT-PCR assay for *Mx1* specific mRNA in infected animal lungs infected was developed. In parallel, infections were performed in BALB/c mice which have a non-functional *Mx1* gene due to a large frameshift deletion^26^. As seen in figure 3A, treatment with ΔNS1 resulted in the early induction (24 hours post infection) of *Mx1* specific mRNA in both A2G and BALB/c mice. In contrast a very faint band was present in A2G mice infected with hvPR8 virus at the same time post infection and no specific mRNA was detected in mock infected mRNA.

**Figure 3.**
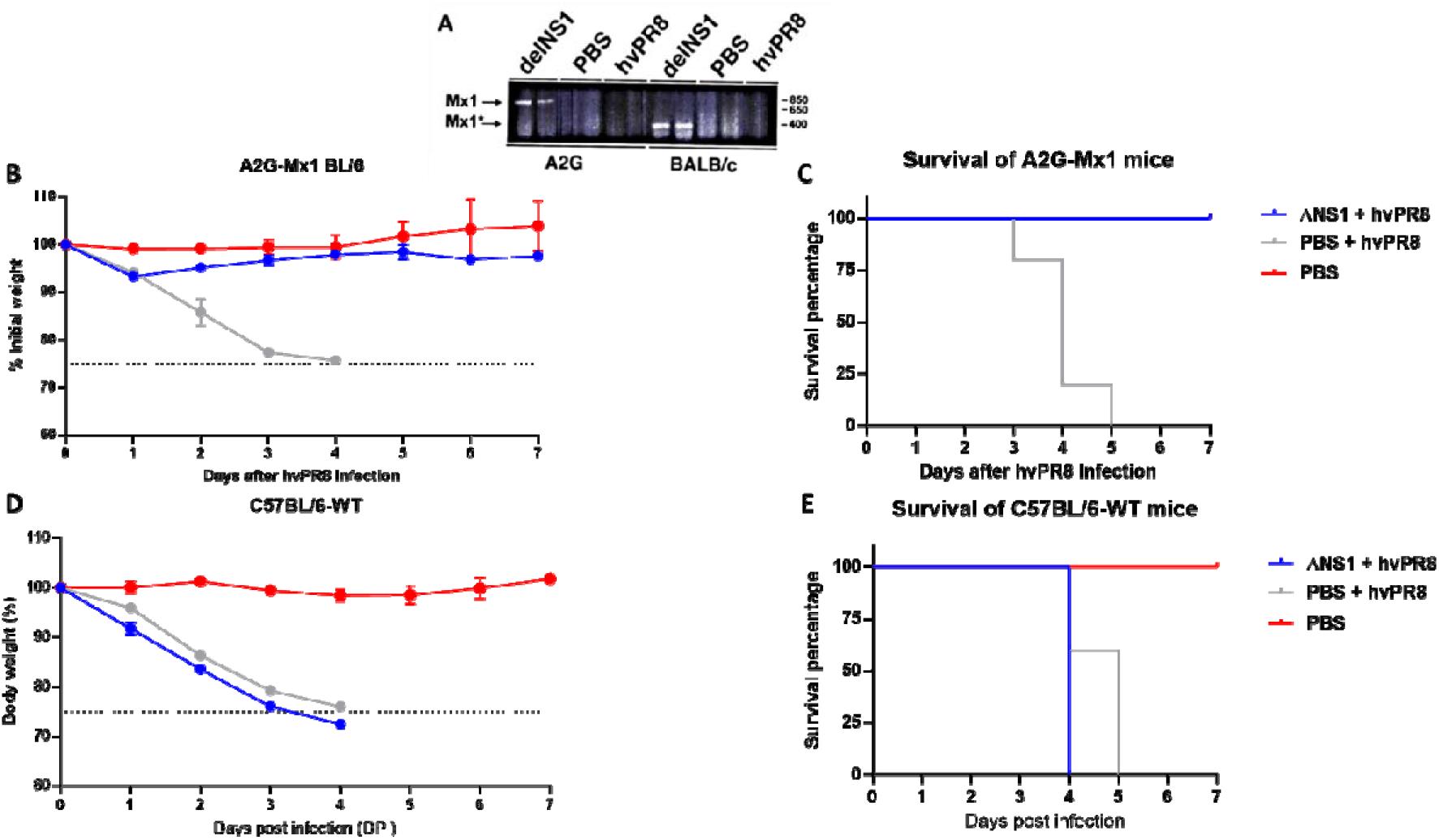
Dose dependent pre-treatment of ΔNS1 protects A2G-M×1 mice but not wild-type C57BL/6 from a lethal hvPR8 virus challenge. **(A) Induction of *Mx1* specific mRNA expression in ΔNS1 virus infected mice**. Groups of two A2G or BALB/c mice were intranasally treated with PBS or 2.5×10^5^ PFU of ΔNS1 hvPR8 influenza viruses. 24 hours post challenge, total RNA present in lung tissues were extracted and were used for RT-PCR reactions using *Mx1* specific primers. PCR products were run in an agarose gel; the arrows indicate the predicted size of amplified cDNA from *Mx1* genes pf A2G mice (M×1) and BALB/c mice (M×1*).**(B,C,D,E)** Sex matched 6 weeks old groups C57BL/6-A2G-M×1 mice or C57BL/6-wild-type mice were either intranasally pre-treated with PR8-ΔNS1 (5×10^6^ PFU; n=5 per group), sterile PBS (n=5) 12 hours before a lethal challenge of hvPR8 (5×10^5^ PFU; n=5) or treated with only sterile PBS (n=2). **(B)** Morbidity of C57Bl/6-A2G-M×1 mice. **(C)**. Survival of C57Bl/6-A2G-M×1 mice. **(D)**. Morbidity of C57Bl/6-wild-type mice. **(E)**. Survival of C57Bl/6-6-wild-type mice.

### ΔNS1 mediated protection from hvPR8 is *Mx1*-mediated

As the M×1 protein is one of the most potent IFN inducible gene products with anti-influenza virus activity in mice, it is quite possible that the ΔNS1-mediated protection seen in A2G mice is M×1-mediated. To test this hypothesis, we compared the antiviral activity of ΔNS1 in A2G mice and in C57BL/6 mice. C57BL/6 mice harbour a non-functional *Mx1* gene due to a known deletion^26^ and were used as a back-cross genetic platform for the original A2G strain to generate the M×1 positive A2G mice used in our experiments. A dose of PR8-ΔNS1 containing 5×10^6^ PFU given 12H before a lethal hvPR8 challenge protected all A2G-M×1 mice (n=5) in both morbidity and mortality in comparison to the PBS pre-treated group (n=5). However, all five M×1-deficient mice in the wild-type C57BL/6 group that were given the same dose of PR8-ΔNS1 succumbed to death by a lethal hvPR8 challenge. The morbidity data for these mice based on body weight was also consistent with lack of protection after ΔNS1 treatment from hvPR8 challenge, indicating that the antiviral effect on IAV induced in mice by ΔNS1 treatment is dependent on the IFN-inducible gene *Mx1* w (Figure 3D and 3E).

### ΔNS1 viral treatment inhibits the replication of hvPR8 virus in A2G mice lungs

To better understand the ability of the ΔNS1 virus to inhibit replication of the hvPR8 virus in the lungs, A2G mice were intranasally treated with 2×10^5^ PFU of ΔNS1 virus alone, 2×10^4^ PFU of hvPR8 alone or treatment of 2×10^5^ PFU of ΔNS1 virus 24 hours before infecting them with 2×10^4^ PFU of hvPR8 virus. Mice were sacrificed at three- and six-days post infection and the lung homogenates were titrated in MDCK or Vero cells (Supplementary table.2). A reduction of hvPR8 titers in lungs by fourfold was observed when mice were pre-treated with ΔNS1 virus. Furthermore, mice solely infected with ΔNS1 virus had titers below the detection limit (<10 PFUml^−1^), while not showing any significant reduction of bodyweight. It was apparent that infection by hvPR8 virus without ΔNS1 administration resulted in the increase of lung weight by a factor of two or three in comparison to mice that were pre-treated with ΔNS1 virus. In the context of this study, increased lung weights are suggestive of lymphocytic infiltration and pulmonary disease during Influenza virus infection^28, 29^.

### Attenuated influenza viruses via a mutation in the Neuraminidase (NA) gene does not confer ΔNS1-like antiviral properties

Antiviral properties observed thus far in this study is from an attenuated influenza virus lacking the NS1 gene (ΔNS1). To confirm that the protective effects observed here are not due to the attenuation caused by the lack of a gene but specifically due to the lack of NS1, the antiviral property of ΔNS1 virus was compared to that of a the recombinant D2 influenza virus. The D2 virus contains a base-pair mutation in the dsRNA region formed by the non-coding sequences of its NA gene. This mutation is responsible for a 10-fold reduction in the NA protein levels as well as a one-log reduction in viral titers within a multicycle growth curve^30^. The latter D2 strain has also been shown to be highly attenuated in mice with a LD_50_ of more than 10^6^ PFU upon intranasal administration^31^. Identical doses (2.5×10^5^ PFU) of D2 or ΔNS1 viruses were intranasally administered to A2G mice four hours prior to challenge with 5×10^6^ PFU of hvPR8. Although a prolonged survival was seen in one of the animals who received D2, pre-treatment with D2 was ineffective in protecting A2G mice from hvPR98 virus-induced disease and death (Figure 4).

**Figure 4.**
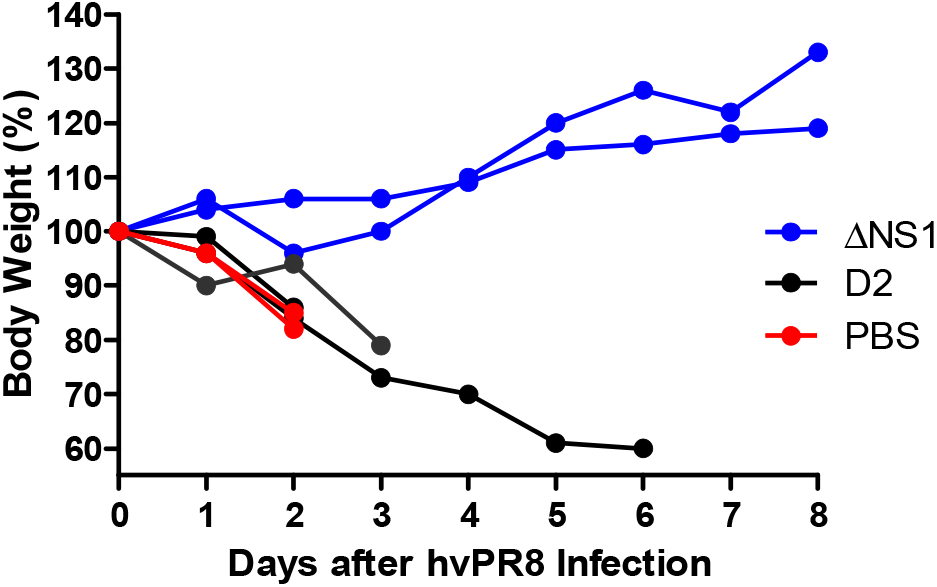
Comparison of the antiviral properties in A2G mice of recombinant influenza A viruses ΔNS1 and D2. A2G mice were intranasally treated with PBS or 2.5×10^5^ PFU of ΔNS1 or D2 viruses for 24 hours before infection with 5×10^6^ PFU of hvPR8 influenza virus. Bodyweight changes and survival were monitored. Data shown are from individual mice.

### ΔNS1 viral treatment prevents death by Sendai virus (SeV) in C57BL/6 mice

Given the fact, that the antiviral effects against hvPR8 mediated by ΔNS1 viral are facilitated by an IFN mediated mechanism (*Mx1* gene induction), we speculated that ΔNS1 treatment should protect mice from infections by other IFN sensitive viruses. Sendai virus was used in this study due to its pneumotropic nature and sensitivity to IFN in *Mx1* deficient mice^32, 33^. As seen in Figure 1B, treatment with ΔNS1 inhibited Sendai viral replication in embryonated chicken eggs. Moreover, upon two intranasal administrations of 2.5×10^5^ PFU of ΔNS1 virus to C57BL/6 mice at times −24 and +24 hours or −8 and +72 hours, mice infected with 5×10^5^ PFU of Sendai virus were protected from death (Figure 5A). The C57BL/6 mice used here are *M×1^−/−^* and it is indicative that the mouse nuclear M×1 protein does not have any antiviral activities against cytoplasmic viruses such as Sendai virus^34^. The efficacy of ΔNS1 treatment was compared against three doses of IFN-β using the Sendai virus challenge model. Treatment with the highest dose of IFN-β (2×10^5^ U) protected mice from death induced by Sendai virus comparable to treatment with 2.5×10^5^ PFU of ΔNS1 virus (Figure 5B).

**Figure 5.**
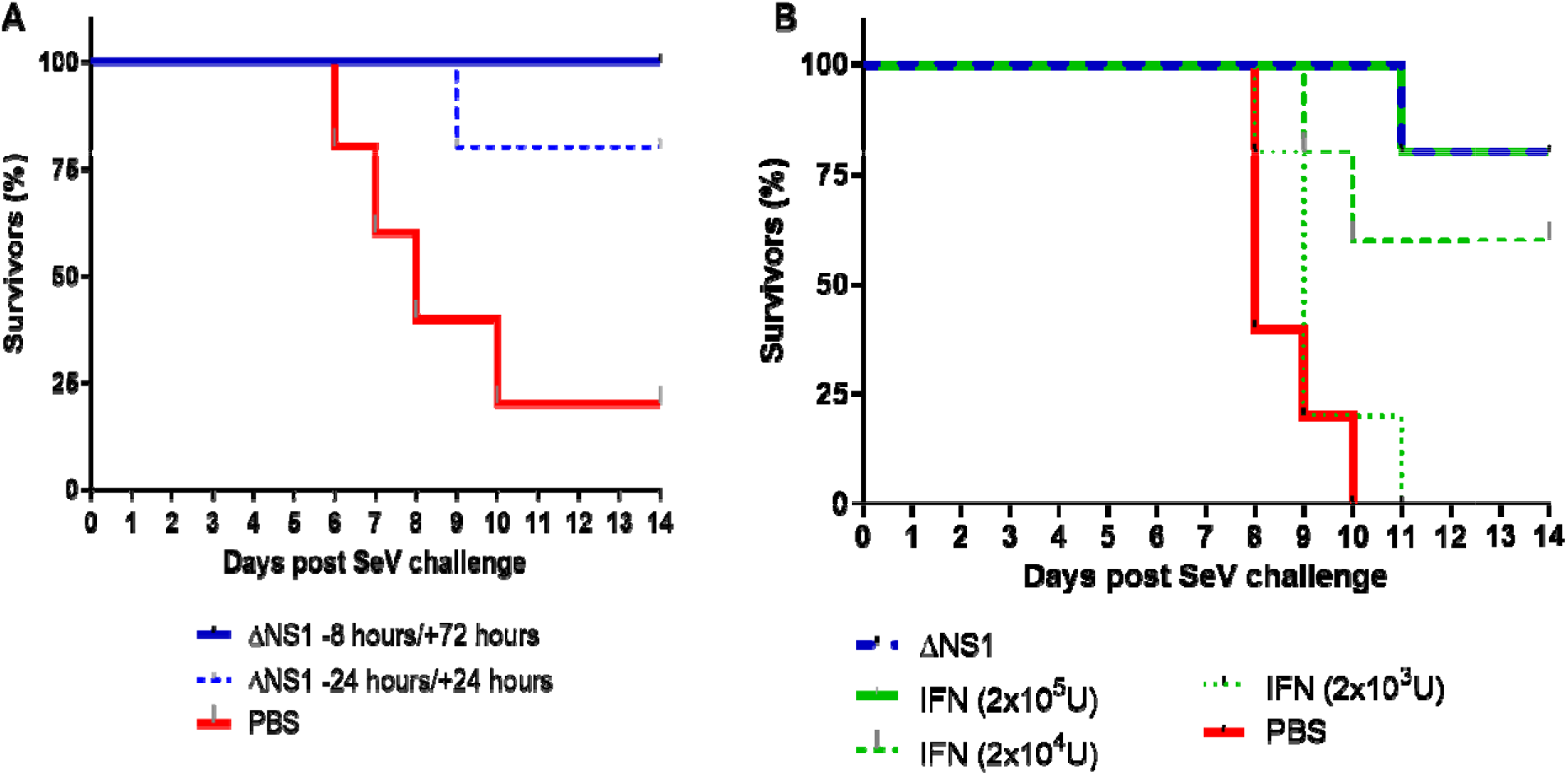
Treatment with ΔNS1 influenza virus protects C57BL/6 mice against lethal infection with Sendai virus. All mice were challenged intranasally with a lethal dose of Sendai virus corresponding to **(A)** 5×10^5^ PFU or **(B)** 1.5×10^5^ PFU. The percentage of mice surviving the challenge is represented. **(A)** Groups of five mice were treated intranasally with 2.5×10^5^ PFU of ΔNS1 virus at the indicated times. **(B)** Groups of five mice were intranasally treated at −24h and +24h with respect to the infection with Sendai virus with 2.5×10^5^ PFU of ΔNS1 or with the indicated amounts of IFN-β.

### ΔNS1 virus treatment inhibits viral replication of SARS-CoV-2 virus in K18-hACE2-C57Bl/6 murine lungs

Given the emergence of the devastating COVID-19 pandemic, we assessed whetherprophylactic treatment with ΔNS1 would hinder the replication of SARS-CoV-2. We used the transgenic mouse model that supports the replication of SARS-CoV2. As controls, we used universal IFN, and SeV defective RNA (SDI) which were previously shown to have an IFN inducing effect. Weight determination in all the treated groups showed no major loss in bodyweight, only one mouse each from the SDI treated group (day 8) and the uIFN treated group (day 12) reached below 75% bodyweight (Figure 6A). Deaths (4 out of 5) in the mock treated group occurred between days 6-8 post infection. The SDI-RNA treated group lost 2 out of 5 animals on day 8 and 9 while the uIFN group lost one animal out of 5 at a later time point (D12; Figure 6B). While both treatments resulted in reduction of viral titers day 3 and 5 post infection, mice that received ΔNS1 showed significant inhibition of SARS-CoV2 titers in lung homogenates and no detectable infectious viruses at day 5 post infection (Figure 6C).

**Figure 6.**
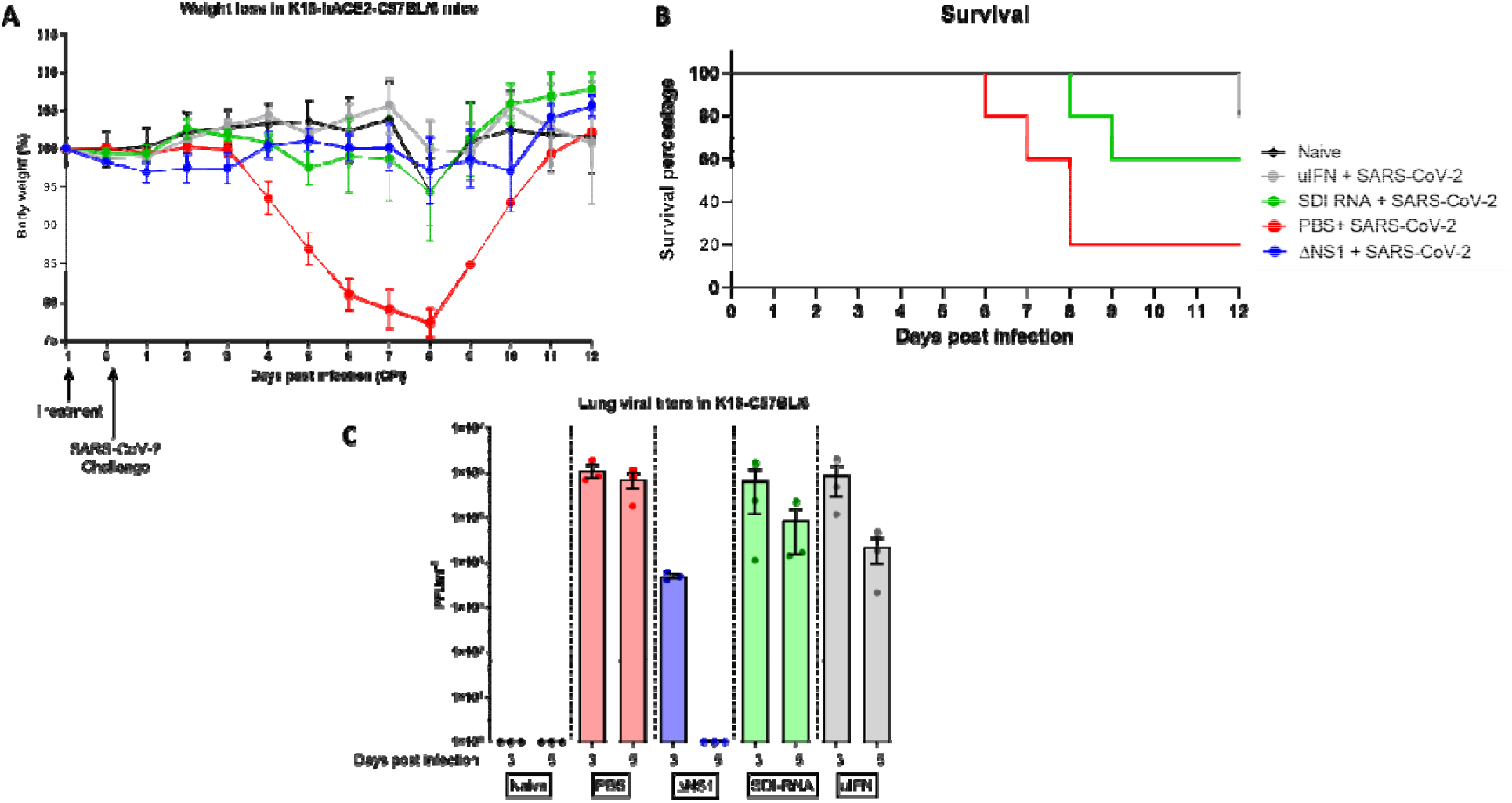
Treatment with ΔNS1 influenza virus inhibits viral replication in the lungs of K18-hACE2 mice challenged with SARS-CoV-2. Mice were intranasally treated with 30 ul containing PBS, 2.5×10^6^ PFU of ΔNS1, 1 μg defective interfering RNA from Sendai virus (SDI-RNA), 2.5×10^5^ U of universal-interferon (uIFN) 24 hours before intranasal challenge with 10^4^ PFU of SARS-CoV-2/USA/WA1 isolate. **(A)** weight-loss was monitored in mice (n=11 for treated groups and n=6 naïve) and **(B)** survival was monitored for 12 days. **(C)** Lungs were harvested at days three and five post infection (n=3 per group per day) were homogenized and were titered in Vero-E6 cells using standard plaque assays.

## Discussion

The NS1 protein of the influenza A virus has been shown to possess IFN antagonist activity whereby it is able to dampen the host innate immune response to provide a favourable environment for the virus to replicate. It has been demonstrated to be highly expressed in the host cytoplasm and nucleus upon viral infection, interacting with a plethora of host factors to inhibit the interferon response^35^. Data show the ability of NS1 to compete with innate immune sensors such as RLR to bind to dsRNA to avoid innate immune detection^36^. Additionally, NS1 has been shown to interact with other innate immune signalling components such as PKR^37^, TRIM25^38^ and CPSF^16^, resulting in lowering of the IFN mediated innate immunity^39^. For these reasons, influenza viruses with impaired NS1 function (and an increased innate immune response) have been under consideration for live attenuated influenza vaccines. There is an existing swine influenza vaccine based on NS1-deficient live attenuated viruses^40^, and clinical trials in humans using an intranasally administered live attenuated ΔNS1 virus have demonstrated potent immunogenicity and good safety profiles. Experimental evidence in mice indicates that the high IFN-inducing properties of ΔNS1 viruses are responsible for their superior immunogenicity as live vaccines^41, 42^.

As ΔNS1 viruses are great IFN inducers, we reasoned that they might provide with innate protection against respiratory virus infection even before the development for an influenza virus specific adaptive immune response. Treatment with ΔNS1 virus inhibited the replication of both homologous and heterologous viruses in eggs (Figure.1). Using the A2G-M×1 mouse model, we demonstrated that the intranasal administration of the ΔNS1 virus induced an antiviral state, which prevented disease and death by a highly pathogenic influenza A virus (hvPR8) which is otherwise lethal^43^. Infection with ΔNS1 virus but not WT viruses yielded detectable levels of *M×1-*specific mRNA levels in lungs 24 hours post infection (Figure 2). A large body of evidence has indicated that the protective impact of IFN against IAV infection in mice is mainly mediated by the IFN inducible antiviral *Mx1* gene^44–46^. Consistently, we found that M×1 was required for the ΔNS1 mediated protection against lethal hvPR8 challenge by comparing M×1 competent A2G--C57BL/6 mice with M×1 deficient WT-C57BL/6 mice.

Data depicted in Figure.2C show that pre-treatment of A2G mice with ΔNS1 virus up to four days before the challenge with hvPR8 virus was effective in preventing disease. The M×1 protein in mice is known to be stable for several days upon its induction and our observations are consistent with the half-life of the M×1 protein described in mice^47, 48^.

Given the inherently attenuated state of the ΔNS1 viruses, it was necessary to confirm that the antiviral state seen here is due to the specific attenuation of the ΔNS1 segment. We used a virus that is known to be attenuated due to its defective neuraminidase segment (D2 virus expressing a full-length NS1) ^31^ to demonstrate that protection is not just mediated by any attenuated IAV (Figure.4). ΔNS1 treated mice were also protected from lethal infection with an influenza-unrelated pneumotropic Sendai virus, suggesting that the IFN-mediated innate immune response induced by ΔNS1 has broad-antiviral effects, rather than being a pathogen-specific immune response. As anticipated for Sendai virus, the abovementioned protection was not *Mx1* mediated and is most likely due to the activation of other ISGs such as OAS or PKR upon the ΔNS1-mediated IFN production^49^.

The feasibility of ΔNS1 virus as a prophylactic treatment to induce a type I interferon response to prevent acute respiratory infections from IFN sensitive viruses was demonstrated in the current study. Type I interferon administration has been used to treat a range of human diseases ranging from infections such as hepatitis B and C^50,51^ to other non-communicable diseases such as melanomas^52^ and hairy-cell leukaemia^53^. Although IFN has been promoted as a therapeutic agent, administration of exogenous interferon comes with a set of undesirable side effects^54, 55^, arguably due to its causing major endocrine and metabolic changes in the host^56^. Therefore, various groups have attempted alternative ways to induce local type I IFN responses using different strategies. Some of these strategies were topical administration of plasmid DNA coding for IFNα1 in the mouse eye to protect against HSV-1 encephalitis^57^, liposomic intranasal treatment using dsRNA to induce IFN^58^ as well as recombinant viral vectors such as adenoviruses^59^ and hepatitis B viruses to express type I IFN to protect against infection and tumor regression^59^. Despite these experimental attempts to study the efficacy of IFN, it is still unclear whether virally induced IFN is more or less toxic efficient that IFN itself. This indicates that further work is needed to be done to ascertain the suitability of recombinant viruses as IFN inducers for therapeutic purposes. The physiological half-lives and binding affinities of different types of interferons are well studied and their half-lives can range from minutes to several hours, depending on the type of IFN^60^. Our data showed antiviral properties of ΔNS1 virus for up to four days before the viral challenge. While it is known that therapeutic properties and doses of different types of IFNs are highly variable due to their differential effects contributed by the ISGs, most therapeutic properties of type I interferons are yet to be completely understood^61, 62^. In this instance, comparable prophylactic responses were obtained by the administration of either 2×10^5^ U of IFN-β or 2×10^5^ PFU of ΔNS1 virus (Figure.5B). However, it is acknowledged that different subsets of IFN-regulated genes may differ in their relative transcriptional induction between treatments.

We also demonstrated that prophylactic treatment using ΔNS1 significantly inhibited viral replication in a relevant mouse model that can be infected with WT SARS-CoV-2 and is known to result in lethal infection^63^(Figure 6). This agrees with reports that state that SARS-CoV-2 is sensitive to IFN^64^. Interestingly, a similar level of reduction in viral titers was not seen upon intranasal inoculation of universal-IFN nor defective interfering RNA derived from SeV (SDE-RNA; a RIG-I agonist with known adjuvanting properties)^65^. While these treatments resulted in a better outcome in comparison to PBS pre-treatment, high amounts of viral titers were still observed day three and five post infection. Although weight loss and survival were best in the ΔNS1 group, the uIFN treated group showed a protective phenotype indicating that uIFN treatment was better than that provided by SDI-RNA. The difference observed here is likely due to the stimulation of multiple innate immune mechanisms by ΔNS1 which potentially primes cells to confer a broad antiviral phenotype. However, analysis of differentially expressed genes (particularly ISGs) via a technique such as bulk RNAseq would provide more insights in explaining the observed protective effects against COVID-19 in the K18 mouse model.

In conclusion, we report that prophylactic treatment with an attenuated influenza A virus lacking the NS1 gene induces an innate antiviral response which provides protection against IFN-sensitive viruses in both embryonated chicken eggs and mice. These *in vivo* data further validate previous observations showing the IFN-antagonistic properties of the NS1 protein of influenza A viruses^13, 66–68^, while highlighting the role of NS1 in inhibiting IFN induction during influenza A virus infections. We also provide evidence for its therapeutic potential as a prophylactic to protect against acute respiratory infections caused by IFN-sensitive viruses including the causative agent of COVID-19 pandemic. ΔNS1 viruses are being clinically developed as live attenuated influenza virus vaccines and in clinical trials they have shown to induce protective antibodies and no adverse responses in human volunteers^21–23^. Here we show that ΔNS1 viruses have the potential to induce immediate protection against viral infection prior to the induction of specific long-lasting protective adaptive immune responses^69, 70^. Our results should encourage further research on the use of IFN-inducing, live attenuated virus vaccines, to confer innate and adaptive protection against virus pathogens.

## Methods

### Cells and viruses

Recombinant influenza A viruses were generated using reverse genetics as previously described^13, 30^ A derivative of the A/PR/8/34 (PR8) defined as highly virulent PR8 (hvPR8) was kindly provided by O. Haller and J.L. Schulman. Strain 52 of Sendai virus was obtained from the ATCC. Vero cells, Madin-Darby bovine kidney (MDBK) cells, baby hamster kidney (BHK) cells or embryonated chicken eggs were used to propagate the following viruses as per standard protocols; Influenza A ΔNS1, hvPR8, PR8, A/WSN/33, A/X-31/H3N2, Influenza B/Lee/40, Sendai virus and vesicular stomatitis virus (VSV). Madin-Darby canine kidney (MDCK) cells or Vero cells were plated to obtain confluent monolayers and plaque assays were performed as previously described and an agar overlay in DMEM-F12 including 1 μgml^−1^ of trypsin was used. MDCK, cVero and BHK cells were cultured in DMEM in the presence of 10% FBS and penicillin-streptomycin. The chicken embryo fibroblasts (CEF) purchased from ATCC was maintained in MEM as suggested by ATCC. Vero-E6 cells (ATCC® CRL-1586™, clone E6) were grown in DMEM containing 10% FBS, non-essential amino acids, HEPES and penicillin-streptomycin. SARS-CoV-2, isolate USA-WA1/2020 (BEI resources; NR-52281) was handled under BSL-3 containment in accordance with the biosafety protocols validated by the Icahn School of Medicine at Mount Sinai. Viral stocks were amplified in Vero-E6 cells in the above media containing 2% FBS for three days and were validated by whole-genome sequencing using the Oxford-MinION platform.

### Animal studies

All animals used in the study were used at 6-10 weeks of age. A2G mice were kindly provided by Dr. Heinz Arnheiter while the BALB/c and C56BL/6 mice were purchased from Taconic Farms. Hemizygous female K18-hACE2 mice on the C57BL/6J genetic background (Jax strain 034860), were used to conduct studies with SARS-CoV-2 in BSL3 conditions. Anesthetized animals were intranasally infected using 30 to 50 μl of appropriately diluted viruses or PBS containing the indicated amounts of recombinant murine IFN-β (Calbiochem), universal-IFN (PBL assay science) SDI-RNA^65^. Afterwards, the animals were monitored daily for changes in body weight. All animal studies were done in accordance with the NIH guidelines as well as the guidelines devised by the Icahn School of Medicine with regards to the care and use of laboratory animals.

### Measurement of Interferon

Ten day old embryonated eggs were infected with 10^3^ PFU in100 μl containing either ΔNS1, PR8 viruses or PBS as mock. Next, the eggs were incubated at 37°C and the allantoic fluids were extracted 18 hours post infection. Viral inactivation of the allantoic fluids were conducted by dialysis against 0.1 M KCL-HCL buffer at pH 2 for two days at 4°C. Later, the pH of the samples was adjusted to pH 7 by subsequent dialysis against Hank’s balanced sodium salt solution with 20 mM NA_3_PO_4_ for two more days as described previously^71^. The amount of IFN was titrated according to its ability to inhibit the growth of VSV^72^. In summary, CEF cells in 96wells were treated with 100 μl of different dilutions of the respective samples in tissue culture media. Upon incubating for an hour at 37°C, 200 TCID_50_ of VSV in 10 μl were added to the wells before incubating at 37°C until complete lysis of untreated control cells was observed (approximately two days). As a standard control, recombinant chicken IFN donated by Drs. Peter Staeheli and Bernd Kaspers was used^73^.

### Lung Titration

Four A2G mice were intranasally challenged with 2×10^5^ PFU of ΔNS1 at day −1. During day 0 mice were intranasally challenged with 2×10^4^ PFU of hvPR8 virus. Alternatively, two other groups of four A2G mice were challenged with 2×10^5^ PFU ΔNS1 or 2×10^4^ PFU of hvPR8. Three days post infection, two animals from each group were humanely sacrificed while the rest of the animals were humanely sacrificed six days post infection. Lungs were weighed and homogenized in 2 ml of PBS. Resulting homogenates were clarified via centrifugation at 3000 rpm for 15 minutes at 4°C and the acquired supernatants were tittered by plaque assays using MDCK or Vero cells. Lung homogenates derived from SARS-CoV-2 infected K18 mice were handled and titered in Vero-E6 cells as described previously^74^.

### Detection of *Mx1* Specific mRNA in infected cells

A2G and BALB/c mice were intranasally challenged with 10^5^ PFU of either ΔNS1 or hvPR8 or PBS. Afterwards, lungs were extracted 24 hours post infection, snap frozen, homogenized, total RNA was extracted using TRIreagent (Sigma-Alderich). One microgram of total lung RNA was used to perform a RT reaction in a total volume of 20 μl using *Mx1* specific primer. Two μl of the resulting RT product was used for PCR amplification using *Mx1* specific primers under the following conditions (20 seconds at 95°C, 30 seconds at 55°C, 30 seconds at 72°C for a total of 25 cycles). The sense and antisense primer sequences are as follows; 5’-CAGGACATCCAAGAGCAGCTGAGCCTCACT-3’ and 5’-GCAGTAGACAATCTGTTCCATCTGGAAGTG-3’. The PCR products were analysed using a 1.2% agarose gel. Correct size for the PCR products in A2G mice was 756 bp while it was 333 bp in BALB/c mice due to a deletion in the *Mx1* gene between nucleotides 1120-1543^31^.

## Acknowledgements

The authors acknowledge members of AG-S and PP laboratories for their critical discussions. Technical assistance for the study was provided by Louis Ngyenvu and Richard Cadagan. We thank R. Albert for support with the BSL3 facility and procedures at the Icahn School of Medicine at Mount Sinai, New York. A2G mice were kindly provided by Dr. Heinz Arnheiter (NIH). hvPR8 virus was a generous gift from Drs. Otto Haller (University of Freiburg) and Jerome L. Schulman (Icahn School of Medicine). Recombinant Chicken IFN was a gift from Drs. Peter Staeheli (University of Freiburg) and Bernd Kaspers (University of Munich). This work was partly supported by the Center for Research for Influenza Pathogenesis, a Center of Excellence for Influenza Research and Surveillance supported by the National Institute of Allergy and Infectious Diseases (contract number HHSN272201400008C), by the NIAID funded Collaborative Influenza Vaccine Innovation Centers (contract number 75N93019C00051), by NIAID grants R01AI141226, R01AI145870 and P01AI097092, by DARPA grant HR0011-19-2-0020, and by the generous support of the JPB Foundation, the Open Philanthropy Project (research grant 2020-215611 (5384)), and anonymous donors to AG-S and PP.

## Author contributions

AG-S, PP, RR, MS and TM conceived the project. RR, MS, HZ, TK and SJ conducted experiments while MS, RR analysed the data and wrote the manuscript.

---

**Supplementary Table 1.** Units of IFN present in the allantoic fluid of 10-days embryonated chicken eggs which were inoculated with WT A/PR8/34 or ΔNS1 influenza A viruses.

| Virus<sup>a</sup> | Egg number | IFN (Uml<sup>-1</sup>)<sup>b</sup> |
| --- | --- | --- |
| **rWT PR8** | 1 | <16 |
|  | 2 | <16 |
| **ΔNS1** | 3 | 400 |
|  | 4 | 400 |
| **Mock** | 5 | <16 |
|  | 6 | <16 |

<sup>a</sup>Eggs were inoculated with 10<sup>3</sup> PFU of rWT-PR8 or ΔNS1 virus
<sup>b</sup>Amount of IFN in the allantoic fluid was measured 18 hours post inoculation

**Supplementary Table 2.**
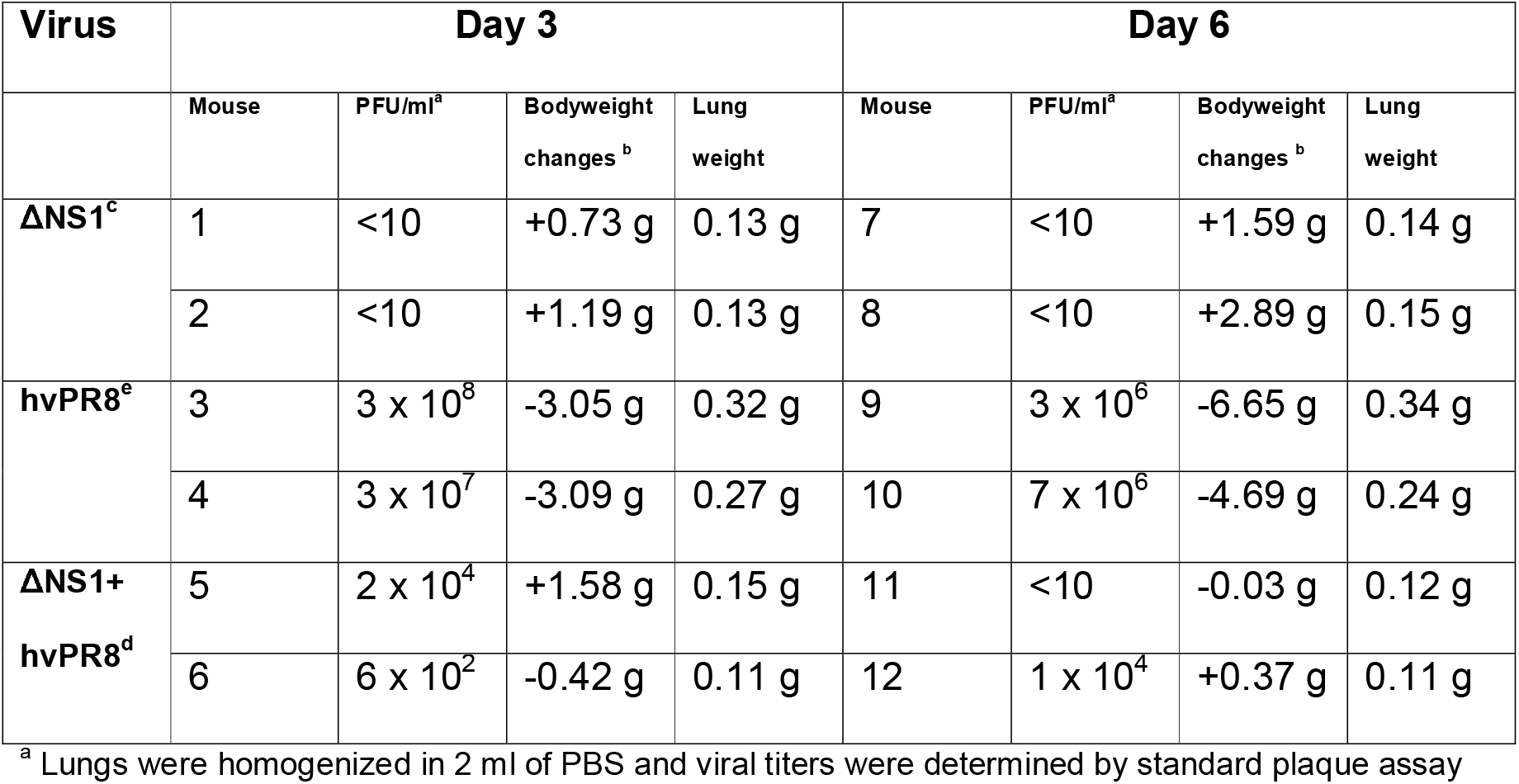
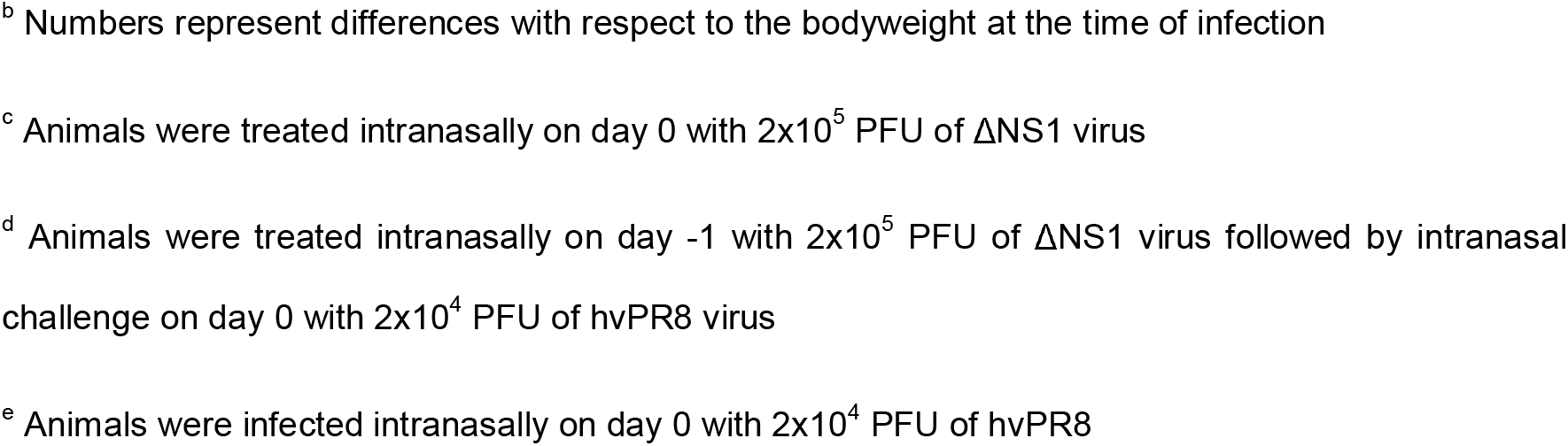
Viral titers, bodyweight changes and lung weights in A2G mice infected with ΔNS1 and hvPR8 viruses.

| Virus | Day 3 |  |  |  | Day 6 |  |  |  |
| --- | --- | --- | --- | --- | --- | --- | --- | --- |
|  | Mouse | PFU/ml<sup>a</sup> | Bodyweight changes<sup>b</sup> | Lung weight | Mouse | PFU/ml<sup>a</sup> | Bodyweight changes<sup>b</sup> | Lung weight |
| **ΔNS1<sup>c</sup>** | 1 | <10 | +0.73 g | 0.13 g | 7 | <10 | +1.59 g | 0.14 g |
|  | 2 | <10 | +1.19 g | 0.13 g | 8 | <10 | +2.89 g | 0.15 g |
| **hvPR8<sup>e</sup>** | 3 | 3 x 10<sup>8</sup> | -3.05 g | 0.32 g | 9 | 3 x 10<sup>6</sup> | -6.65 g | 0.34 g |
|  | 4 | 3 x 10<sup>7</sup> | -3.09 g | 0.27 g | 10 | 7 x 10<sup>6</sup> | -4.69 g | 0.24 g |
| **ΔNS1+ hvPR8<sup>d</sup>** | 5 | 2 x 10<sup>4</sup> | +1.58 g | 0.15 g | 11 | <10 | -0.03 g | 0.12 g |
|  | 6 | 6 x 10<sup>2</sup> | -0.42 g | 0.11 g | 12 | 1 x 10<sup>4</sup> | +0.37 g | 0.11 g |

<sup>a</sup> Lungs were homogenized in 2 ml of PBS and viral titers were determined by standard plaque assay
<sup>b</sup> Numbers represent differences with respect to the bodyweight at the time of infection
<sup>c</sup> Animals were treated intranasally on day 0 with $2 \times 10^5$ PFU of ΔNS1 virus
<sup>d</sup> Animals were treated intranasally on day -1 with $2 \times 10^5$ PFU of ΔNS1 virus followed by intranasal challenge on day 0 with $2 \times 10^4$ PFU of hvPR8 virus
<sup>e</sup> Animals were infected intranasally on day 0 with $2 \times 10^4$ PFU of hvPR8

